# From storms to warming seas: a long-term metabarcoding survey in port communities unveils high genetic diversity and ecological resilience of non-indigenous species

**DOI:** 10.64898/2025.12.20.695700

**Authors:** J. Zarcero, A. Antich, O.S. Wangensteen, M. Rius, X. Turon

## Abstract

Ports are key gateways for the introduction and spread of non-indigenous species (NIS), yet the ecological and genetic temporal dynamics of these introductions remain poorly understood. Long term temporal monitoring is essential to unravel invasion processes, anticipate biodiversity shifts, and inform effective management and biosecurity strategies. In this study, we conducted a five-year (2019-2024) monthly metabarcoding survey in a northwestern Mediterranean port using artificial collectors. By sequencing a fragment of the Cytochrome Oxidase I gene, we identified 2,225 Molecular Operational Taxonomic Units (MOTUs), including 93 NIS. In addition, we examined both interspecific temporal patterns and intraspecific trends of genetic diversity over time. Although NIS accounted for only 4% of total species richness, they represented over 26% of total read abundance, underscoring their strong influence on community structure. Interestingly, NIS had a significantly more homogeneous species composition through time than native species. In 2020, the passage of the Gloria storm reshaped community dynamics, triggering a temporary rise in species richness and MOTU counts, likely due to an influx of native taxa and a gradual decline and increased NIS dominance. Genetic analyses revealed that NIS exhibited higher haplotypic diversity and lower genetic differentiation than native taxa, suggesting sustained gene flow, likely facilitated by maritime transport. MOTUs with longer temporal persistence, particularly among NIS, also showed greater intraspecific diversity, supporting the “insurance hypothesis” and highlighting the role of genetic variability in resilience and invasion success. Overall, our findings showed that NIS, despite their low species richness, maintain high abundances, connectivity, and genetic diversity over time. These attributes likely enhance the NIS ability to persist in dynamic and disturbed port environments, and provide key information for understanding the invasion process. This study highlights the need to integrate genetic diversity metrics into marine biomonitoring assessments and management.

## 1 Introduction

The spread of non-indigenous species (NIS) is considered one of the most serious threats to the preservation of biodiversity worldwide (Guy-Haim et al., 2018; Hulme et al., 2009; Kumschick et al., 2015; Molnar et al., 2008; Pyšek & Richardson 2010; Tempesti et al., 2020). In line with this, the European Union’s Marine Strategy Framework Directive (EU, 2008) has designated NIS as a crucial indicator of environmental health, requiring systematic data collection on their population size, geographic distribution, and ecological impacts (EU, 2017). Effective management of NIS in marine environments hinges on rigorous monitoring in space and time (Gittenberger et al., 2023). Temporal monitoring of NIS, in particular, is a neglected field (Haubrock & Soto, 2023), yet it is essential to capture seasonal and long-term dynamics, as well as NIS responses to environmental changes. Without this temporal framework, monitoring efforts risk missing critical invasion stages and/or range expansion patterns, undermining the efficiency of biosecurity responses.

The Mediterranean Sea is recognised as one of the world’s major marine biodiversity hotspots (Coll et al., 2010; Myers et al., 1990) and, at the same time, has become one of the seas most threatened by NIS due to both the high number of introduced species (Katsanevakis et al., 2014; Ulman et al., 2017; Zenetos 2018; Zenetos et al., 2022) and the rapid pace at which species introductions occur (Zenetos et al., 2022). These species thrive in ports and marinas, where they benefit from conditions such as lower native species diversity, warmer temperatures, higher nutrient inputs, and a greater availability of substrates for attachment, compared to the open sea (Railkin, 2004). Mediterranean NIS have important impacts, not only on native biodiversity, but also on ecosystem services and human health (Rotter et al., 2020; Tsirintanis et al., 2022). The globalisation of maritime transport has further accelerated NIS translocation, acting as a key vector through multiple pathways, such as biofouling on ships and ballast water (Hulme et al., 2009; Kolar & Lodge, 2002; Tempesti et al., 2020). In this context, NIS introduced via ports not only cause ecological imbalances but also have significant socio-economic impacts on the maritime sector, as they damage infrastructure and require substantial financial resources for maintenance (Roy et al., 2023). Port areas are thus considered strategic points for the monitoring of NIS arrivals throughout the Mediterranean (Occhipinti-Ambrogi & Galil, 2010; Romeo et al., 2015; Ulman et al., 2019).

To carry out proper biosecurity surveillance of port and marina communities, genetic metabarcoding techniques have been proven to be an effective and fast way of unveiling biodiversity patterns and detect NIS (Duarte et al., 2021; Rey et al., 2020). Additionally, metabarcoding enables the identification of early life history stages, such as larvae and recruits, which are often the phases during which marine NIS are transported, and it also has the potential to discriminate among cryptic species. As a result, metabarcoding techniques have been increasingly used in recent years to survey port communities, particularly with a focus on NIS (e.g., Ammon et al., 2018; Holman et al., 2019; Lavrador et al., 2024; Zarcero et al., 2024). However, most studies to date have relied on snapshot or short-term surveys (usually covering seasonal variation) conducted at single or multiple ports (e.g., Rey et al., 2020; Azevedo et al., 2020). While some long-term temporal studies have been carried out using metabarcoding on open coast environments (e.g., Daraghmeh et al., 2025; Sildever et al., 2023, Turon et al., 2025), on port communities these temporally-oriented studies are currently lacking. Yet they may provide invaluable information about the dynamics underlying NIS spread and persistence.

Metabarcoding data can also be used to assess intraspecies genetic patterns through the analysis of DNA sequence composition of the inferred molecular operational units (i.e., metaphylogeography, see Antich et al., 2021; Turon et al., 2020). This approach allows the integration of community-level diversity studies with analyses of community genetic diversity and connectivity patterns (Antich et al., 2023, Overcast et al 2023, Riley et al 2025). This framework is highly relevant in the context of NIS research, since repeated NIS introductions can increase their genetic richness while simultaneously promoting human-mediated genetic homogenisation (Hudson et al., 2022; Roman & Darling 2007; Zarcero et al., 2025). Examining the genetic diversity patterns of NIS provides insight into their invasive potential and the role they might play in displacing native species (Stepien et al., 2002). Likewise, the levels of gene flow following the establishment of NIS can determine a population’s success or failure in colonising new territories, its resilience to environmental stress, and the outcome of competitive interactions with other organisms (Wade et al., 2017). Ports constitute ideal experimental field laboratories for assessing ecological and evolutionary changes in response to entirely new connectivity networks and environmental conditions (Touchard et al., 2023, Lilli et al., 2025). Long-term studies are essential for a proper monitoring of NIS dynamics in these areas. To analyse these evolutionary mechanisms, temporal data are essential (Touchard et al., 2024).

Here, we conducted a long term temporal analysis of marine communities in a port of the northwestern Mediterranean, with a particular focus on NIS and their changes in diversity and genetic composition. Specifically, we tested the following hypotheses: (1) NIS exhibit low richness but high relative abundances, reflecting their colonisation success. (2) NIS have higher haplotypic diversity and greater temporal homogenisation than native species, as a result of recurrent introductions mediated by shipping traffic (David 2018), while native species experience greater genetic isolation and are more dependent on local processes (Blanco González et al., 2016; Hall & Kuhlmann 2013; Voisin et al., 2005). (3) Species with higher haplotypic diversity show greater temporal persistence, particularly among NIS, consistent with classical ecological theory and the insurance hypothesis, which links genetic variability with stability and adaptive potential (Frankham 2005; Hughes et al., 2008; Quigley 2022; Reed & Frankham, 2003; Yachi & Loreau, 1999). Finally, we also took advantage of an extreme climatic event occurring during our monitoring (Gloria storm in 2020) that allowed us to test the resilience of NIS after disturbances.

## 2 Methods

### 2.1 Field sampling and sample processing

Recently developed artificial collectors (so-called “POMPOMs”, Zarcero et al., 2024) have been proven effective at capturing both early life-history stages and DNA shed by adults, as well as particulate organic matter carrying DNA molecules, thus providing a comprehensive view of the biodiversity present. Four replicated collectors were placed monthly in the port of Blanes (41.674425 °N, 2.799687 °E), located on the Catalan coast of the Northwestern Mediterranean Sea. This port features 1,486 linear meters of docks and moorings, encompassing fishing and recreational activities. The “POMPOM” collectors were deployed from February 2019 to February 2024, with monthly replacements (note that during the first cold-season replacement was done bimonthly, resulting in 10 temporal sampling points in 2019 and 12 in subsequent years). Three collectors from each month and location were selected for metabarcoding analysis, while the fourth served as a backup. The cleaning protocol for the collectors followed the guidelines from (Zarcero et al., 2024), using sterilised 10 cm nylon brushes. Temperature was recorded hourly throughout the study period using HOBO[Pendant data loggers placed next to the collectors, and daily averages are presented in Fig S1.

During the timeline of the monitoring, in January 2020, Storm Gloria hit the Northwestern Mediterranean (Amores et al., 2020). This was an extreme event without precedent in the area (Pérez-Gómez et al., 2021). The storm surge and wind waves up to 8 m altered marine communities. Specifically, at the port of Blanes the main dyke was broken and seawater flooded the port through this opening and also overflowed the breakwaters.

### 2.2 DNA extractions and sequencing

DNA extractions were carried out using the DNeasy PowerMax Soil Kit (Qiagen). For each sample, 5 grams of homogenised material obtained from each collector were used. A fragment of 313 bp (modal length) of the Cytochrome Oxidase 1 (COI) gene was then amplified using the generalist Leray-XT primer set (Wangensteen et al., 2018), designed for eukaryotes. This set includes the forward primer jgHCO2198: 5’-TAIACYTCIGGRTGICCRAARAAYCA-3’ (Geller et al., 2013) and the reverse primer mlCOIintF-XT: 5’-GGWACWRGWTGRACWITITAYCCYCC-3’ (Wangensteen et al., 2018).

Both primers were tagged with an 8-base sequence tag. The same tag was used for both the forward and reverse primers of each sample to minimize the effect of inter-sample chimeras, and tags differed by at least 3 bases. To increase sequence diversity and improve Illumina base calling, between two and four degenerate bases (N) were incorporated before the tags on both primers. The amplification was carried out using AmpliTaq Gold 360 Master Mix (Applied BioSystems) supplemented with Bovine Serum Albumin, as described in (Wangensteen et al., 2018). The PCR procedure consisted of an initial denaturation step at 95 °C for 10 minutes, followed by 45 cycles of denaturation at 94 °C for 60 seconds, hybridization at 45 °C for 60 seconds, and elongation at 72 °C for 60 seconds, followed by a final elongation step of 5 minutes. The amplification products were purified and concentrated using the MinElute PCR Purification Kit (Qiagen), and positive results were confirmed via gel electrophoresis. Negative samples were processed using sand charred in a muffle furnace, and some PCR blanks with no DNA template were included for each library generated (for a total of 23 PCR blanks and 21 negative controls). Libraries were prepared using the BIOO NEXTFLEX PCR-Free DNA-Seq Kit (Perkin-Elmer) and sequenced on a partial Illumina NovaSeq lane with 2 x 250 bp paired-end sequencing at Novogene Company. All procedures were performed in a sterilised laminar flow cabinet, with UV light activated between samples to avoid contamination.

### 2.3 Bioinformatic workflow

Bioinformatic analyses were performed using a pipeline based on Obitools3 software (Boyer et al., 2016), employing bash and R 4.0.2 scripts. Briefly, paired-end reads were aligned using illuminapairedend, retaining only those with an alignment quality score greater than 40. The reads were then demultiplexed using ngsfilter, which removed sequences with unmatched primer tags and trimmed the primer sequences. The obigrep and obiuniq functions were applied to filter by length and to pool identical sequences. As the fragment amplified was 313 bp long in most eukaryotes, we retained sequences 310, 313, 316, and 319 bp to keep codon integrity. Chimeric amplicons were removed using the Uchime_denovo algorithm from VSEARCH v2.7.1 (Rognes et al., 2016). The sequences were then denoised using the DnoisE program (Antich et al., 2022), a modification of the Unoise algorithm (Edgar, 2016), which accounts for natural variability in the three codon positions of coding genes. Denoising was performed within samples with an alpha parameter of 4, and an auto-computed entropy to generate Exact Sequence Variants (ESVs) (Antich et al., 2021). Several filters were applied to the ESV dataset: 1) ESVs with over 10% of reads found in blanks or negative controls were removed as they likely represent contaminations. 2) A dual abundance filtering was applied, removing (i) ESVs representing less than 0.005% of total reads within each sample, and (ii) ESVs with fewer than 5 reads across samples after the first filtering step.

The ESV dataset was then clustered into molecular operational taxonomic units (MOTUs) using SWARM v3.1.3 with a distance parameter (d) of 13, following Antich et al., (2021). SWARM is a fast, threshold-free algorithm that initially connects all reads with a distance less than d, and then refines clusters based on topological criteria related to their internal abundance structures (Mahé et al., 2014). The most abundant ESV in each MOTU was selected as the representative sequence. MOTU assignment information for each ESV was obtained from the SWARM output and added to the ESV dataset.

Taxonomic assignment of the MOTU representative sequences was performed using the mkLTG program (Meglécz, 2024) with the COInr database (Meglécz, 2023). The default parameters of the mkLTG procedure were adjusted as described in Zarcero et al., (2025). To avoid overclassification (Mugnai et al., 2023), Insecta and Arachnida were excluded from the reference database, while maintaining orders that include marine mites, using the select_taxa script of the mkCOInr program (Meglécz, 2023). Only MOTUs assigned to Metazoa were retained for further analysis. Taxonomic assignments were checked manually to remove human and other non-marine MOTUs (likely contaminants). The final dataset refinement involved the application of the LULU post-clustering correction procedure (Frøslev et al., 2017). This step combines similarity and co-occurrence metrics to detect and correct erroneous MOTUs by pooling reads with the correct ones. A modified version of the LULU function (https://jesuszarcero/LULU_corrected) was used to fix a previously detected bug (https://github.com/tobiasgf/lulu/issues/8).The ESV dataset was updated accordingly.

Finally, ESVs were checked for erroneous sequences, including nuclear mitochondrial inserts (NUMTs), following Turon et al., (2020) and Zarcero et al., (2024). All ESVs whose sequences contained codon stops were removed after verifying the 12 metazoan mitochondrial genetic codes from the Biostrings R package (Pagès, 2017). Sequences with changes in any of the five conserved amino acids across metazoans in the studied fragment (Pentinsaari et al., 2016) were also excluded. MOTUs for which all ESVs were eliminated were deleted from the final MOTU table.

Non-indigenous species detection was performed using the NCBI-BLAST algorithm (Korf et al., 2003) and the NISdb v3 database (Zarcero et al., 2024, https://github.com/jesuszarcero/NISdb), which includes curated sequences of introduced species in the Mediterranean. This database was compiled from the available literature on NIS in the Mediterranean. COI-5P sequences for these species were obtained from the Barcode of Life Data System (BOLD, https://www.boldsystems.org/). If sequences were not available in BOLD, the NCBI database was searched and species identification was cross-verified with taxonomic literature. For each NIS, sequences were collapsed into unique haplotypes. A thorough manual curation was performed to ensure data accuracy, involving extensive BLAST searches and removal of potentially erroneous sequences when BLAST results indicated a mismatch or conflicting species assignment. This version of NISdb represents an update to the original database created by Zarcero et al., (2024). BLAST results were filtered to retain assignments with at least 97% identity and 90% sequence coverage. These assignments were then compared with the corresponding MOTU assignments from the mkLTG procedure to verify the accuracy of NIS detection.

### 2.4 Interspecific analyses

Downstream analyses were conducted on three distinct datasets, depending on the specific question addressed. One of them comprised the complete set of detected MOTUs (ALL MOTUs dataset). Note that this dataset includes numerous MOTUs that were not identified at the species level, making it impossible to determine whether they are NIS or native. We also considered separately the NIS dataset (comprising MOTUs assigned to NIS through the above procedure). Finally, to enable more targeted comparisons with the NIS dataset, we selected the MOTUs that were identified at the species level and could be classified as native (as checked with the relevant literature), forming the NAT dataset. Analyses have been performed using as temporal units both months and years. For the latter, we did not follow calendar years but instead used 12-month periods from February to February to align with the sampling schedule.

Analyses were conducted with the ‘vegan’ R package (R Core Team, 2023, Oksanen et al., 2019), and plots were created with the ‘ggplot2’ R package (Wickham, 2011), unless otherwise stated. Rarefaction curves and species accumulation curves were generated with the functions *rarecurve* and *specaccum*, respectively. The MOTU richness and Shannon diversity values were calculated following rarefaction to the minimum number of reads across all samples, using the *rarefy* function. These results were organised into monthly and yearly groups. The resulting metrics were compared across categories with ANOVAs and post hoc Tukey tests. To assess the taxonomic composition of the samples, MOTUs were grouped into major metazoan phyla, and bar charts were generated to display the average composition per season in terms of proportion of reads and proportion of MOTUs for the different datasets. Phyla representing less than 5% of the reads were grouped under "Others." The "Unidentified" category corresponded to metazoan reads that could not be assigned at the phylum level. Upset plots were used to visualise patterns of MOTU sharing across years for the different datasets.

For β-diversity analyses, the relative read abundance of each MOTU in every sample was used without applying rarefaction, and the Bray-Curtis dissimilarity index (BC) was calculated. These values were then employed to create reduced-space representations of the samples through non-metric multidimensional scaling (NMDS) using the *metaMDS* function from the vegan package, with separate analyses for the ALL MOTUs, NIS MOTUs, and NAT MOTUs datasets. Mantel tests (function *mantel*) were used to compare the dissimilarity matrices.

Permutational analyses of variance were performed on the BC matrices for the factors year and season using the PERMANOVA module from the Primer v6 statistical package (Anderson & Walsh, 2013), including pairwise tests for significant factors, with a total of 999 permutations used to generate the statistics. Permdisp tests were also run to check differences in heterogeneity of the data.

The Bray-Curtis dissimilarity data were also analysed at increasing temporal frames by grouping them according to the time-lag between samples, from the minimum (1 month) to the maximum (59 months).

The degree of homogenisation of the samples was examined using the match index (Ahrens et al., 2016):

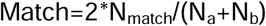

Where N_match_ is the number of shared MOTUs between samples a and b, and N_a_ and N_b_ are the total number of MOTUs in each of them, respectively.

This index quantifies the amount of compositional similarity (MOTU sharing) between samples, and was calculated for the NIS and NAT datasets. The values were compared by paired-sample t-tests.

### 2.5 Intraspecific analyses

Intraspecific genetic diversity values for all MOTUs were obtained as the number of ESVs (proxy for haplotypes) assigned to each MOTU, and an ANOVA test was performed to compare these values between the NIS and NAT datasets. Temporal trends of genetic diversity were also ascertained for these datasets.

A regression analysis was performed between intraspecific diversity and relative abundance values of the MOTUs to assess potential relationships between the two factors, categorizing MOTUs into groups based on their temporal persistence in the samples. To disentangle the effects of both variables on MOTU survival, we performed a stepwise multiple regression analysis to model persistence of each MOTU (in years) as a function of mean relative abundance and mean number of ESVs in the samples where the MOTU was present. The analysis was run with the MASS R package (Venables and Ripley, 2002) in forward mode (adding variables sequentially) using the Akaike Information Criterion (AIC) to evaluate the fitness gain as variables are added to the model. This analysis was performed separately for NIS and NAT MOTUs.

We also computed a genetic differentiation measure among pairs of samples using the D distance (Jost, 2008) with the *pairwise_D* function of the mmod R package (Winter, 2012). This was done for the NIS and NAT datasets. For each pairwise comparison we selected the shared MOTUs with some genetic variation (i.e., at least two ESVs in the two samples being compared). We used as an approximation to haplotype abundances a semi-quantitative index ranging from 0 to 10 (Turon et al., 2025). This index was computed using the relative abundance of each ESV in its corresponding MOTU in the sample: an ESV absent in the sample had a 0 index. An ESV with relative abundance in its MOTU between 0 and 0.1 was assigned a value of 1, and so on. The resulting matrices of genetic distances for NIS and NAT MOTUs were compared among them and with the corresponding Bray-Curtis matrices with Mantel tests (function *mantel*), to check the coherence of the patterns of β-diversity and genetic differentiation. Finally, mean D values of the NIS and NAT datasets were compared with a paired-sample t-test.

Mean genetic distances between pairs of months were obtained by averaging D values of the comparisons between all replicates in these months. These distances were then grouped according to the years of the two months being compared into distances one, two, three, and four years apart. This was done to check changes in genetic differentiation between samples progressively more distant in time.

## 3 Results

### 3.1 Global diversity and datasets

All generated sequences have been uploaded to the NCBI SRA archive (Bioproject PRJNA1268165). After pairing, demultiplexing, quality and length filtering, and chimera removal, we obtained 2,401,982 unique COI sequences. The denoising process resulted in 48,262 ESVs, which were grouped into 6,225 MOTUs. After all filtering steps, retaining only sequences assigned to marine metazoans, our final database (ALL MOTUs dataset) consisted of 156 samples, totaling 2,225 MOTUs, 20,351 ESVs, and 51,366,469 reads.

The final MOTU table, including taxonomic identification and native or NIS status, is provided in Table S1. Additionally, the final ESV table, indicating the MOTU to which each ESV belongs, is presented in Table S2. On average, each sample contained 329,272 ± 21,528 metazoan reads. The mean number of metazoan MOTUs per sample was 123.4 ± 3.16. The sample rarefaction curves (Fig. S2) showed that a plateau was reached in the number of MOTUs per sample, indicating sufficient sequencing depth. However, the MOTU accumulation curves (Fig. S3) did not reach a plateau, suggesting that more MOTUs would be added as additional samples were analysed. Note that this trend is more accentuated in the first two years of study and, particularly, the second one.

We identified 93 MOTUs belonging to NIS using BLAST with the NISdb database, representing 13,383,106 reads and comprising 4,834 ESVs (Table S2). Another 375 MOTUs identified at the species level were classified as native (NAT dataset), accounting for 12,606,473 reads (24.54% of the total) and including 5,276 ESVs (Tables S1 & S2). On average, 4.18% of the MOTUs were assigned to the NIS category, but they represented 26.01% of the reads. At the MOTU level, the proportion of NIS remained more or less stable (Fig. 1A), while a drop in NIS was observed at the read level during the second and third years (Fig. 1B). NAT species represented 16.85% of the MOTUs and 24.54% of the total reads.

**Fig. 1.**
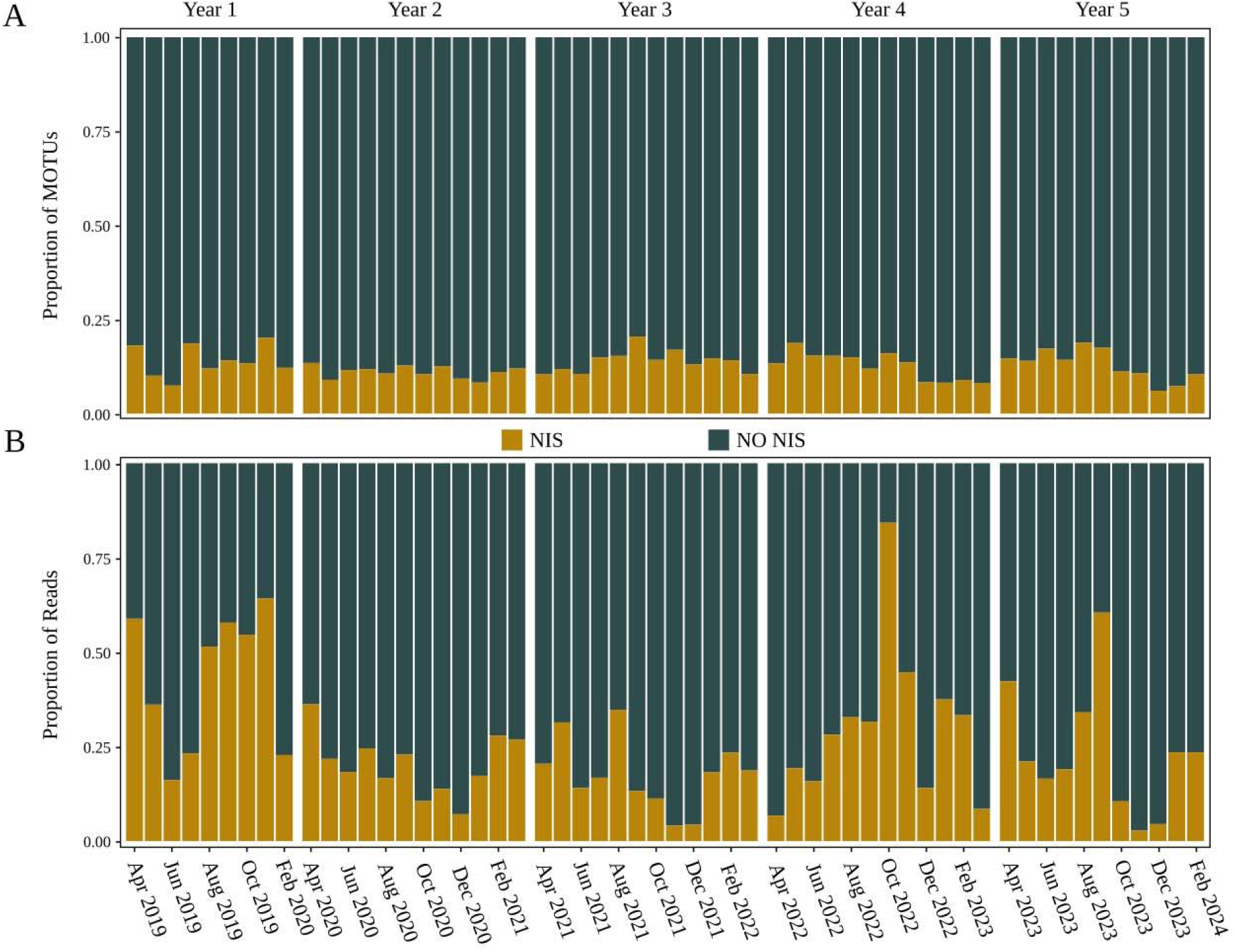
Bar plots showing the proportion of MOTUs (A) and reads (B) corresponding to NIS relative to the rest of the MOTUs for each month over the five-year study period.

### 3.2 Taxonomic composition and patterns

The most abundant group in the ALL MOTUs dataset was cnidarians, with the highest relative read abundance, followed by bryozoans (Fig. S4). The third most abundant group was annelids, followed by chordates, arthropods, molluscs and "Others." The "Unidentified" category accounted for between 3% and 27% of the reads in each season. In the NIS dataset, annelids, cnidarians, and arthropods were, in this order, the dominant phyla in terms of relative read abundance. The temporal variation in relative abundances in the NIS dataset did not show a consistent pattern. Annelids dominated in the first and last years, and partially in the fourth year. For the rest of the time, cnidarians were overall the most abundant in terms of relative reads. In the NAT dataset, cnidarians were the most abundant group during most sampling dates throughout the monitoring period, followed by annelids, molluscs, arthropods, and bryozoans. However, right after Storm Gloria, an abrupt shift in taxonomic composition was observed, with bryozoans becoming the most abundant group in subsequent months (see Fig. S4).

The patterns of shared MOTUs in an upset plot (Fig. S5) showed that many of the MOTUs observed (880 out of 2,225) were exclusive to the second and first years, in that order. Next were the MOTUs that appeared in all years (223), followed by the MOTUs unique to the third, fourth, and fifth years. Conversely, fewer MOTUs were shared across two, three, or four years (between 3 and 94 MOTUs). A similar pattern was observed for the NAT dataset (Fig. S5). However, the NIS showed a clearly different picture, with a substantial proportion of MOTUs (29 out of 93) being present at all years (Fig. S5).

### 3.3 Patterns of **C:**- and **β**-diversity

The mean MOTU richness values, separated by year, are shown in Fig. 2 for the ALL, NIS, and NAT datasets. The corresponding Shannon diversity values are given in Fig. S6. The data were rarefied to the minimum number of reads of any sample, specifically 24,675 reads. The highest mean values for both richness and diversity were observed during the second year, following Storm Gloria, followed by a decrease in the subsequent years. This trend was observed in the three datasets for richness and diversity, except for the diversity of NAT MOTUs, which was higher in the third year. The analysis of variance followed by pairwise tests for the ALL MOTUs dataset showed significant differences in richness values between year 2 and each of the other years, whereas Shannon diversity did not differ significantly among years (Table S3). In the NIS dataset, the analysis of variance revealed significant differences in richness between years 1 and 2 compared with years 4 and 5, while no significant differences among years were detected for Shannon diversity. Similarly, for the NAT MOTUs dataset, the analysis of variance for richness showed significant differences between years 1 and 2, as well as between year 2 and years 4 and 5, whereas Shannon diversity did not differ significantly among years (Table S3).

**Fig. 2.**
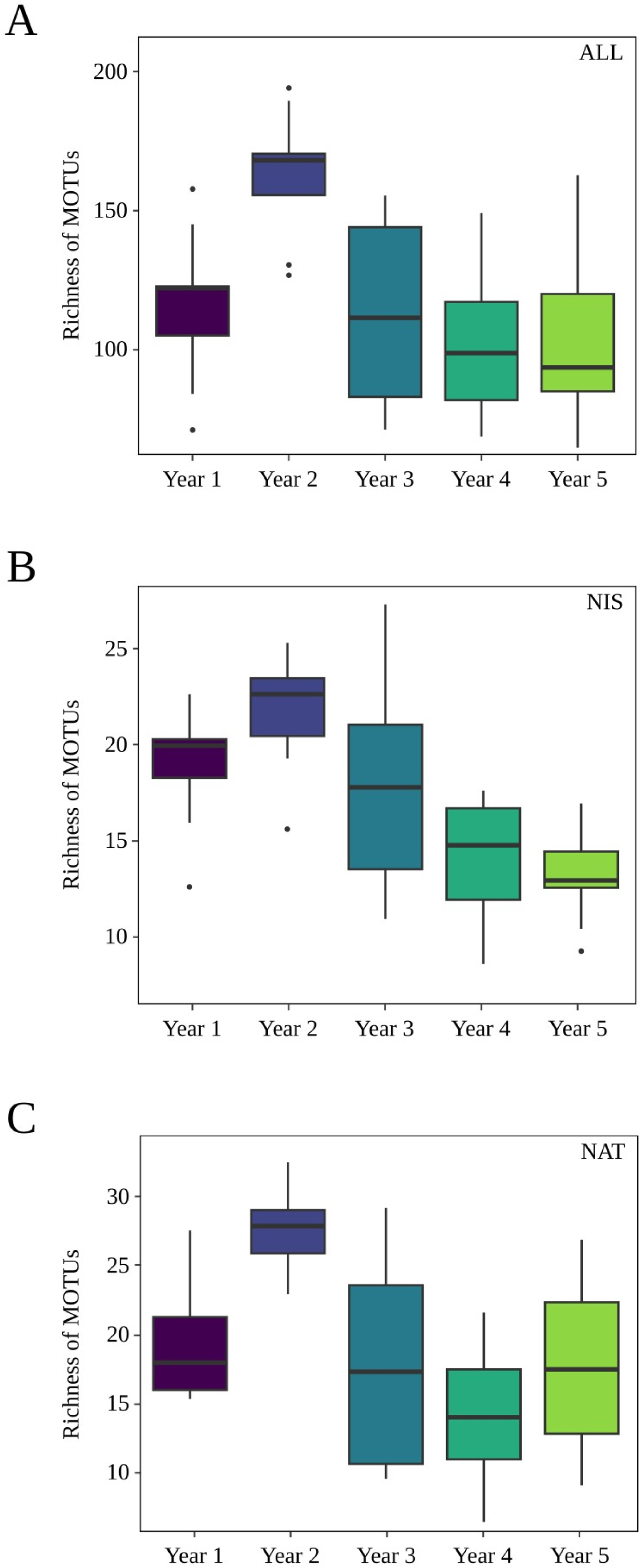
Box plots showing MOTU richness values for each year of the study: the whole dataset (A), NIS dataset (B), and NAT dataset (C). Horizontal lines indicate medians, boxes represent the first and third quartiles, whiskers indicate the 10th and 90th percentiles, and outliers are shown as dots.

When the richness results are plotted by month (Fig. S7), the increase in richness coinciding with Storm Gloria and the following months was apparent. The third year values decreased and remained relatively low for the fourth and fifth year. Again, the patterns were coherent for the three datasets considered. Diversity values (Fig S8) showed a similar pattern, albeit diversity actually decreased for the ALL and NAT MOTUs datasets in February 2020, the month just after Storm Gloria, in spite of a marked increase in MOTU richness. This can be explained by a highly skewed distribution of reads in that month, dominated by a few MOTUs with very high abundances, resulting in a reduction in diversity. The skewness parameter jumped from 5.083 to 14.948 the month before and after the storm, in the ALL MOTUs dataset. Skewness values of the NAT dataset were 2,916 (before) and 7,878 (after).

The NMDS configurations obtained revealed a trend for Summer-Autumn samples to appear separated from the Spring-Winter ones, suggesting a role of seasonality (Fig. 3). There was, however, some overlap, and the separation of seasons is less clear-cut in the NIS and NAT datasets. It must be noted that, in the ALL and NIS configurations, there were three winter samples that appeared clearly displaced to the right of the first axis. They corresponded to the replicates of February 2020, the month after Storm Gloria. The Mantel tests were significant in all cases (all p<0.001) showing an overall congruence in the spatial configurations of the three datasets. However, the Mantel correlation coefficients were noticeably lower when comparing NIS and NAT dissimilarity matrices (r=0.163) than in comparisons of these with the ALL MOTUs one (r=0.601 and r=0.508, respectively).

**Fig. 3.**
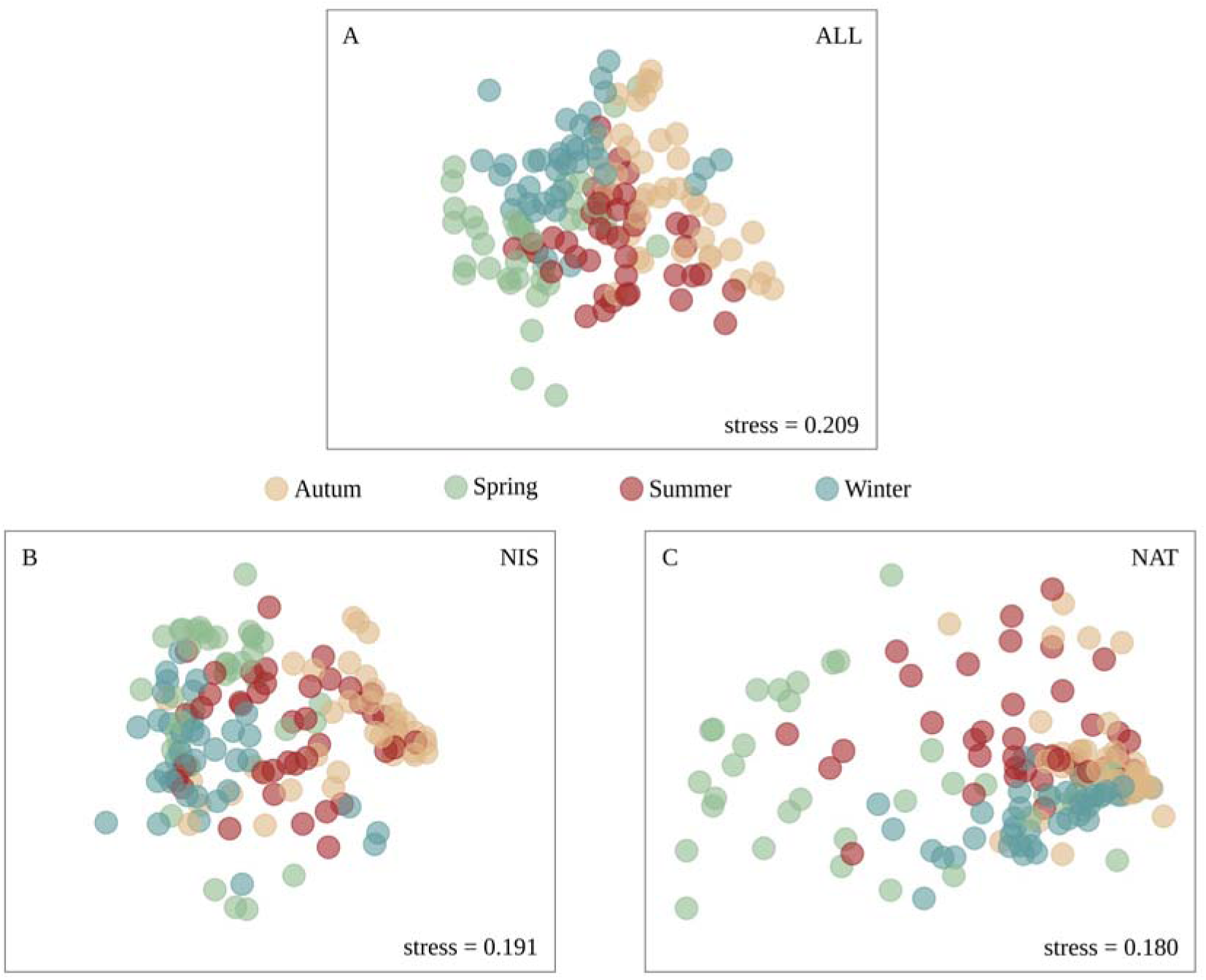
Non-metric MDS configurations for ALL (A), NIS (B), and NAT (C) datasets. Stations are indicated by different colours. Stress values of the final configurations are shown.

The PERMANOVA results showed significant differences in Bray-Curtis dissimilarities according to the factors considered (season, month, and their interaction) for the three datasets (Table S3). Permdisp values were also significant for the two main factors in the NAT dataset, indicating heterogeneity of dispersion in the values for this dataset. In the presence of a significant interaction, pairwise comparisons were performed for levels of one factor at each level of the other factor (Table S3). Overall, these comparisons showed that more significant comparisons among seasons were detected in the fifth year, and fewer in the second year. Likewise, Summer and Spring had in general more significant pairwise comparisons among years than Winter and Autumn. The distribution of BC dissimilarities (ALL dataset) showed that they were higher in comparisons between months in the same year (0.713±0.003, mean±SE) than in comparisons of the same month across years (0.675±0.005), and the difference is significant (Wilcoxon test, p<0.001). Markedly lower values were found among replicates of the same month (0.380±0.013).

A plot of the BC dissimilarities at increasing temporal distances (Fig. 4) showed a clear seasonal pattern, as samples cycled with a 12-month period indicating recurrence in the annual sequence of community composition. Dissimilarities were minimal between samples separated by multiples of 12 months and maximal at lags of 12*n+6 months. This pattern was clearer for the ALL and the NIS MOTUs datasets, while it was somewhat obscured in the NAT dataset, especially at the longest temporal distances (over 40 months). It can be seen that the general trend in the three datasets was one of progressively increasing BC dissimilarities as we examined longer time lags (Fig. 4).

**Fig. 4.**
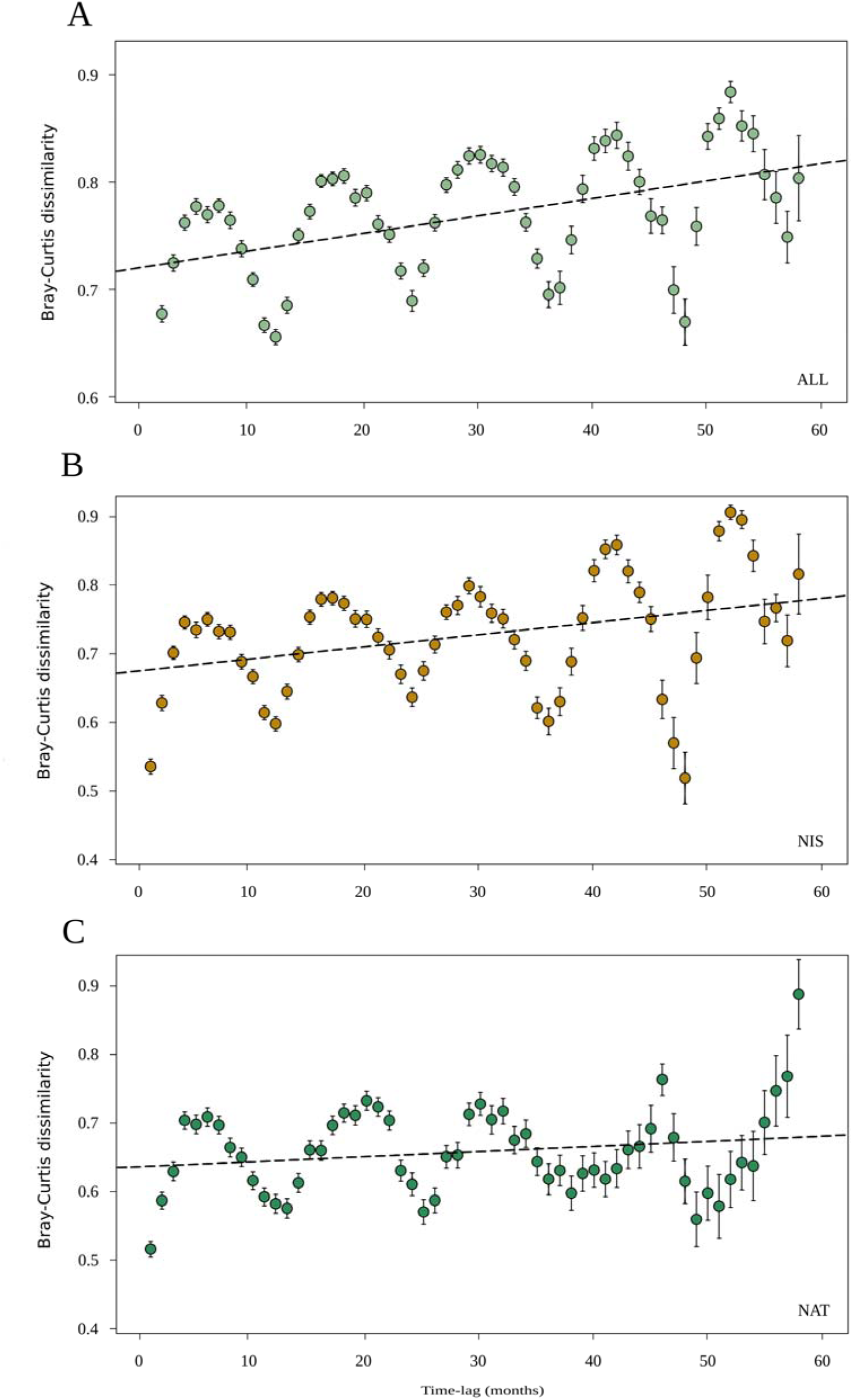
Scatter plot with standard deviation bars for each point showing Bray–Curtis dissimilarities at increasing temporal distances for the ALL (A), NIS (B), and NAT (C) MOTU datasets. The first dissimilarity is calculated between samples one month apart, and the last corresponds to samples 58 months apart.

Finally, the values of the match index across all samples revealed a higher degree of MOTU sharing (i.e., more homogeneity) for the NIS (0.548±0.080) than for the NAT dataset (0.311±0.078), and the difference was significant (paired-sample t-test, p<0.001) (Fig. 5).

**Fig. 5.**
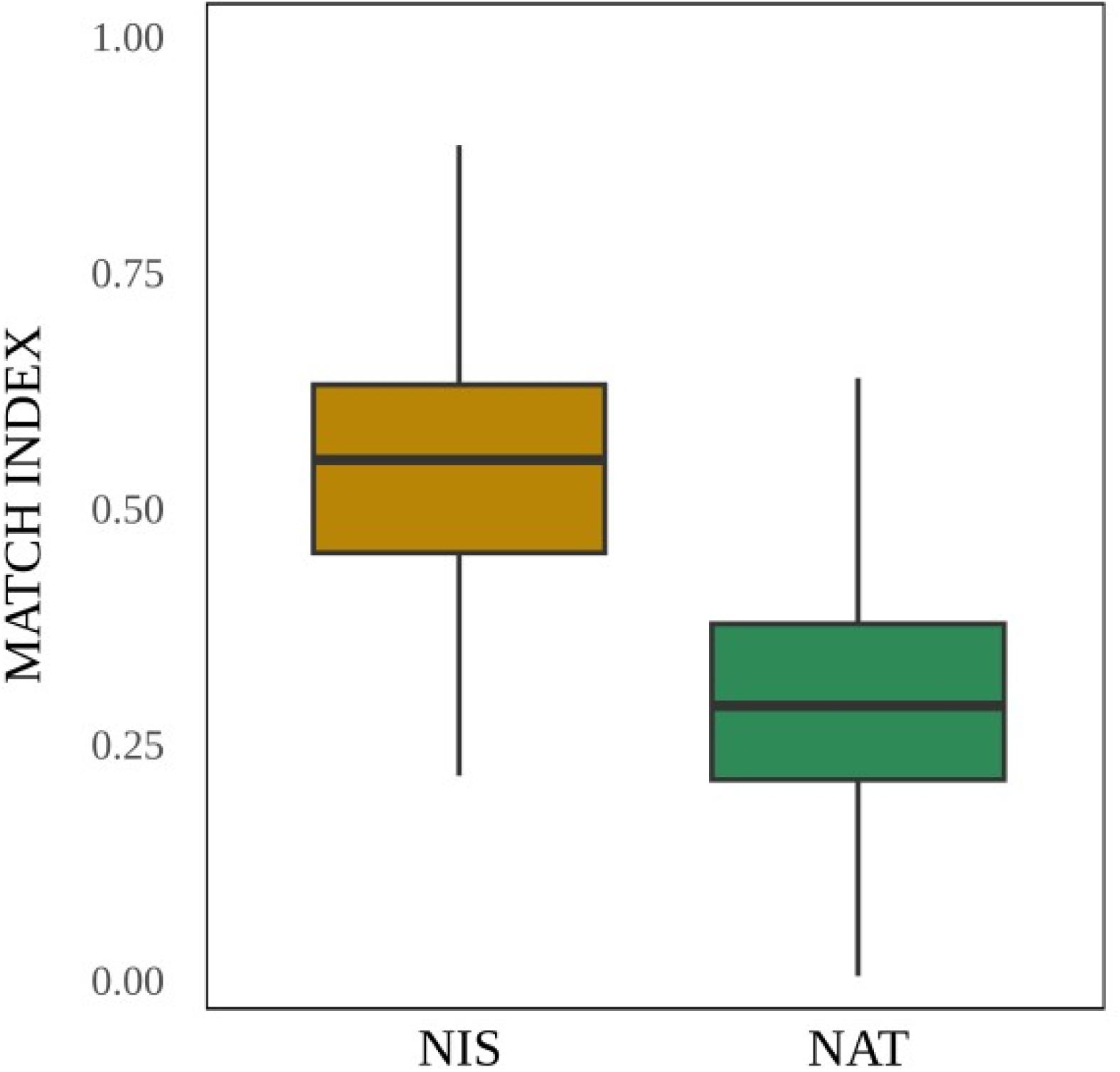
Box plot of Match Index values between samples for NIS and NAT datasets.

### 3.4 Intra-MOTU genetic diversity and differentiation

Overall, the genetic diversity (measured as the number of ESVs per MOTU) was significantly related to the relative abundance for both NIS and NAT MOTUs (NIS, r=0.898, p<0.001; NAT, r=0.647, p<0.001) (Fig. 6, note that one outlier in each dataset with both a high abundance and a high ESV number were eliminated to avoid spurious correlations). In turn, both variables were related to the persistence of MOTUs over time in the samples. MOTUs that were detected during the five years of sampling tended to be high abundance and high genetic diversity MOTUs, while those that persisted only one or a few years were, in general, MOTUs with just one or a few ESVs and with low abundances (Fig. 6). Of the ten MOTUs with the highest genetic diversity, seven also ranked among the ten with the highest read frequencies. All these MOTUs persisted throughout the five years of survey.

**Fig. 6.**
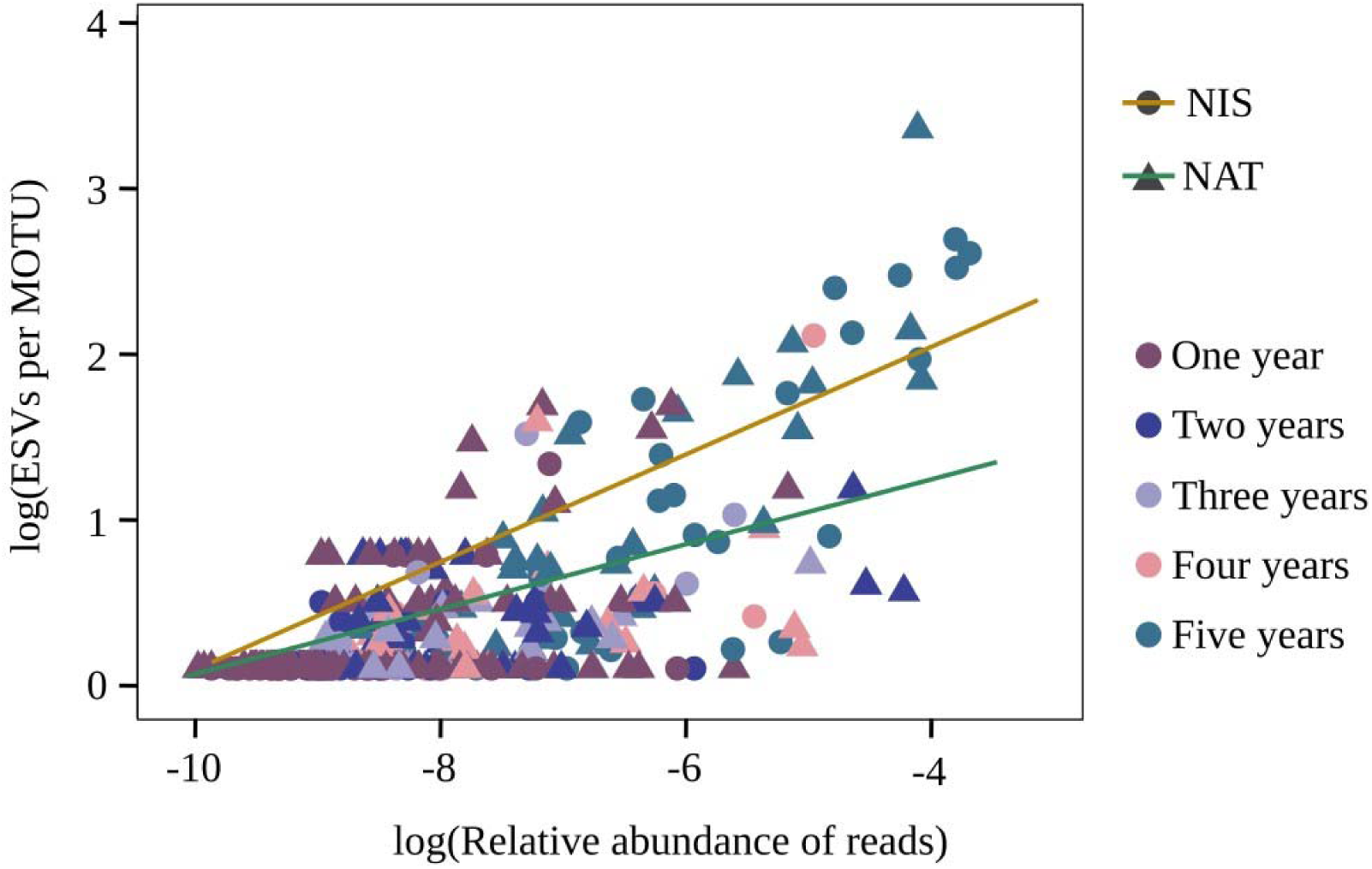
Regression between mean relative read abundance and genetic diversity, calculated as the number of ESVs per MOTU, for each MOTU in the complete dataset. Values were log-transformed to aid visualization. Lines represent trends for NIS and native species, with colours distinguishing different study years.

To separate the effects of both variables, we performed a forward stepwise regression analysis, modelling persistence (in years) as a function of abundance and genetic diversity. For NIS, the variable best explaining survival was the number of ESVs per MOTU and, once the effects of this variable were removed, no significant effect was detected for relative abundance (*r*-squared of the final model = 0.224, p<0.001, Table S3). For NAT, however, both variables were retained in the final model, albeit abundance explained better the variation in persistence times, and the final *r*-squared (0.096, p<0.001,Table S3) was much lower.

The overall values of genetic diversity (ESVs per MOTU) were significantly higher for NIS than for NAT (Fig. 7A, 2.60±0.43 and 1.47±0.12, mean±SE, respectively, t-test p=0.014). In turn, the values of genetic differentiation between months, measured with the parameter D (using the mean of all comparisons among replicates), were significantly lower for NIS than for NAT (Fig. 7B, 0.391±0.001 and 0.443±0.001, mean±SE, respectively, paired-sample t-test p<0.01). There was no significant correlation of D matrices for NIS and NAT (Mantel test, r=0.017, p=0.307), while the D distance (genetic differentiation) showed a low, but significant, correlation with the BC distance (beta-diversity) both for NIS (r=0.183, p<0.001) and NAT (r=0.147, p<0.001) datasets. The genetic differentiation values between months at increasing temporal distances (measured as years apart) did not show significant differences either for NIS or for NAT (Fig. 8, Kruskal-Wallis tests, p=0.087 and 0.397, respectively), indicating no increase of genetic differentiation over time within the studied frame of 5 years.

**Fig. 7.**
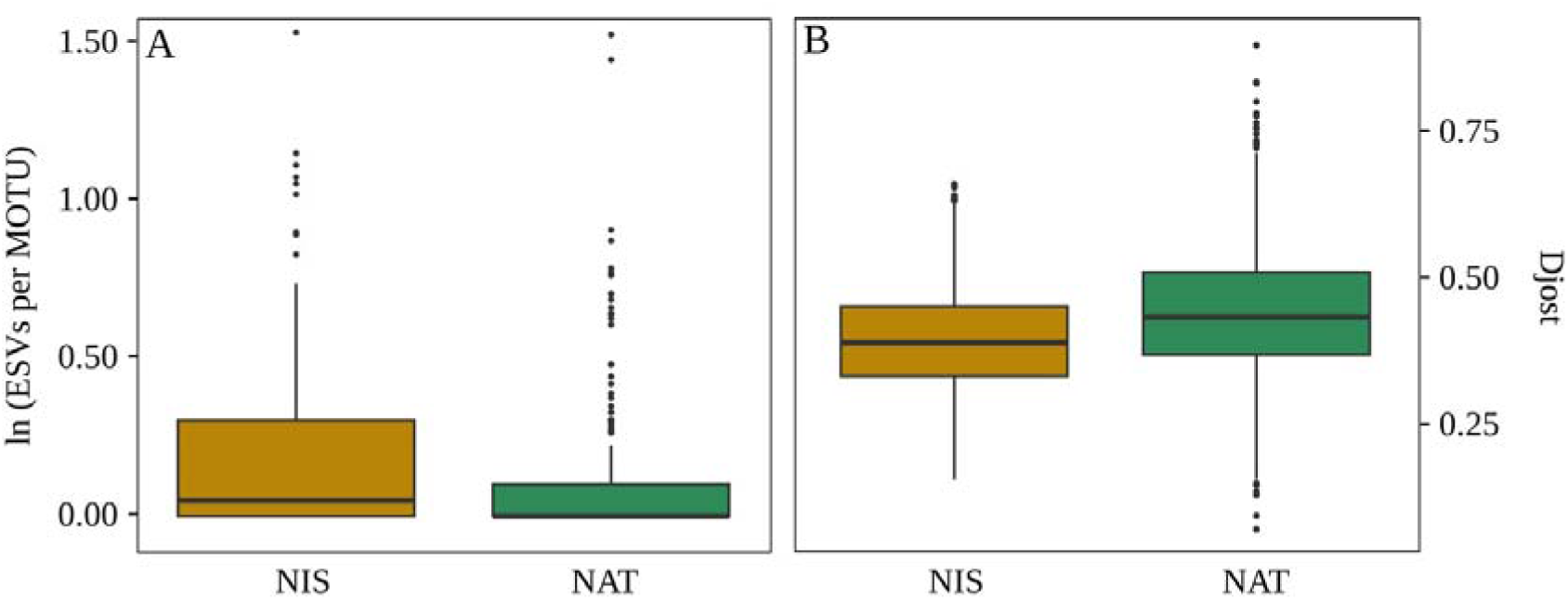
Box plots of the mean proportion of ESVs per MOTU (A) and genetic differentiation with D_Jost_ values (B) for NIS and native species.

**Fig. 8.**
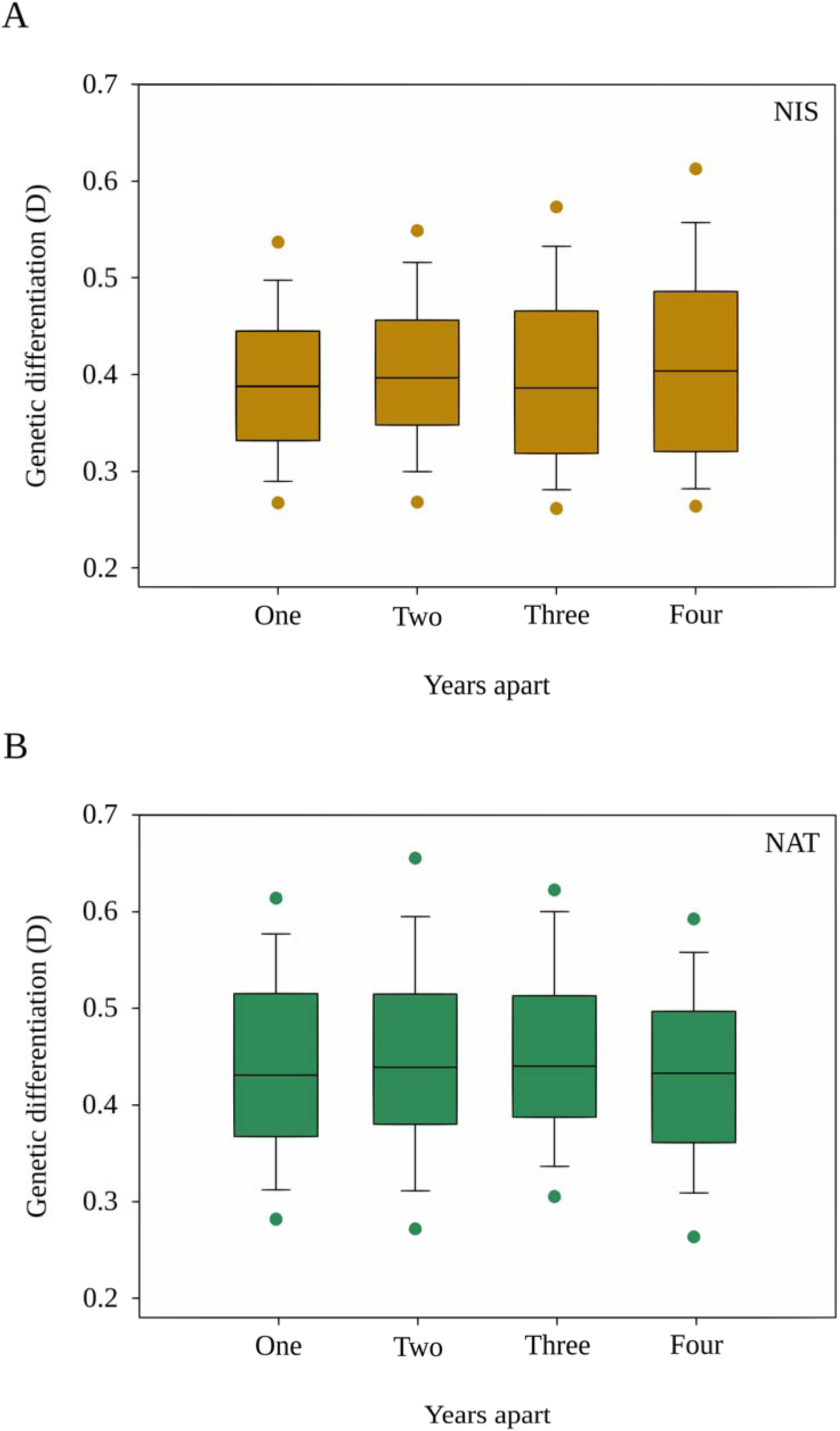
Box plots of genetic differentiation values between months at increasing temporal distances (measured in years) for the NIS (A) and NAT (B) datasets.

## Discussion

Our study unveiled the dynamics of a rich metazoan community in a Mediterranean port with both fishing and recreational activities. Overall, this study provides an integrated understanding of how ecological and genetic processes, together with extreme events, shape the structure and evolution of port metazoan communities, thereby contributing to a better understanding of invasion dynamics and the management of marine biodiversity.

Although MOTUs assigned to NIS constituted a small fraction in terms of richness (ca. 4%), their relative abundance exceeded 26% of the total reads of the community, highlighting a notable influence of these species on community structure and dynamics, as similarly reported in other Mediterranean ports (Ulman et al., 2019; Zenetos et al., 2022). Another 25% of reads could be assigned to verified native species. For both native and NIS, we detected temporal dynamics driven by a seasonal pattern punctuated by extreme events, and ongoing community changes likely associated with the trend of increasing temperatures found.

From the first to the second year of monitoring, there was an increase in the number of MOTUs detected. This finding is coherent with a marked impact of Storm Gloria at the beginning of the second year of study (2020). These findings suggest that the storm event triggered an immediate colonisation process and the arrival of new species, increasing the community’s species richness. During this record-breaking storm (Amores et al., 2020, De Alfonso et al., 2021; Pérez-Gómez et al., 2021), the main breakwater was broken and water also spilled over other breakwaters. This represented an enormous influx of water, probably carrying new species. The sudden increase in richness lasted for another year and diminished again from the third year onwards, returning to seemingly normal levels. However, it must be acknowledged that, as this storm occurred early in our monitoring, we cannot reliably establish a robust baseline status for the port communities. The proportion of NIS numbers (in reads) decreased abruptly after the storm, and remained low for two years, increasing again the 4th and 5th year of study, which likely reflects that the storm-associated influx was mostly of species inhabiting the natural environment with few NIS. The distribution of phyla also showed shifts that can be traced back to the storm event, as the abundance of annelids the first year was replaced by cnidarians over the second and third year (particularly so for NIS). Extreme climatic events are extensively acknowledged as key drivers of biodiversity patterns (Corte et al., 2017; Pagán et al., 2020), and metabarcoding surveys are a powerful method to assess the impact of these alterations (Berry et al., 2019). Furthermore, the increasing frequency and intensity of such events have significant implications for predictive models of species distribution and ecosystem structure (Wernberg et al., 2013).

The analysis of population structure through NMDS, revealed some seasonal differentiation, whereby samples collected during the warmer months (Summer and Autumn) tended to cluster more closely and exhibited higher similarity compared to those collected during colder months (Winter and Spring). A strong seasonal effect was also evident in the temporal variation of dissimilarity values, with periodic peaks occurring every 12 months for the three datasets considered. Notably, dissimilarity values were higher between samples from consecutive months than between samples from the same month across different years, again indicating more pronounced seasonal dynamics compared to interannual variation. Over time, a progressive increase in compositional differences was observed, suggesting long-term structural changes in community composition.

Aside from a clear seasonal pattern, longer-term temperature effects may exist, as peak temperatures were detected in the last two years (2022 and 2023). In these two years, unprecedented heatwaves hit the NW Mediterranean and triggered changes in benthic communities (Boudouresque et al., 2024, Juza et al., 2024, Tejedor et al., 2024, Turon et al., 2025). It is known that climate change is driving marine biodiversity loss worldwide (Penn & Deutsch, 2022), and that historically extreme temperatures favoured the emergence of non-indigenous species (Sorte et al., 2010). In addition, community assembly in anthropised systems is somewhat unpredictable, as communities may include pioneer species and colonisers assembled stochastically (Vieira et al., 2018). . In our case, in the last two years, a decline in richness was observed, particularly among non-indigenous species, that may be due to temperature changes. A longer time frame is likely necessary to disentangle the effects of warming from other factors in the port communities analysed.

Our results showed that NIS presented a distinctly stronger temporal homogenisation in terms of MOTU composition compared with native species, as reflected in a significantly higher match index (ca. 75% higher). Likewise, the values of temporal genetic differentiation were significantly lower for NIS than for native species. In addition, NIS displayed significantly greater haplotypic diversity than the native species present in the port, as also reported by Zarcero et al., (2025). A plausible explanation for these patterns lies in the potential for continuous translocation and gene flow of NIS mediated by maritime traffic, which facilitates the recurrent introduction of species and genetic variants (Roman, 2006; Roman & Darling, 2007; David, 2018). This continuous transport may lead to more homogeneous species composition through time and increased genetic variability of NIS, while simultaneously promoting genetic homogenisation. In contrast, native species appear to be more genetically isolated, with limited exchange among port habitats, which could constrain their haplotypic diversity and restrict their adaptive potential under fluctuating environmental conditions (Blanco Gonzalez et al., 2016; Hall & Kuhlmann 2013; Voisin et al., 2005).

When intraspecific diversity is evaluated in relation to species temporal persistence, a clear pattern emerges, supported by the AIC model: MOTUs with longer temporal persistence in the system show, on average, higher haplotypic diversity and a higher number of relative reads than those with lower persistence, particularly among NIS. These patterns align with classical ecological theory, which posits that greater genetic diversity enhances the adaptive capacity of species, thus increasing the likelihood of long-term survival and persistence (Frankham, 2005; Reed & Frankham, 2003; Torda & Quigley, 2022). According to the “insurance hypothesis”, genetic variability acts as a buffer against environmental disturbances, promoting population resilience and stability (Hughes et al., 2008; Yachi and Loreau 1999). Furthermore, the presence of multiple haplotypes may increase the probability that some genetic variants prove advantageous under changing conditions, providing a foundation for rapid evolutionary responses. In this context, NIS may benefit not only from greater ecological resilience but also from higher genetic diversity, which could support their success in colonising new environments and withstanding major climatic events and other ecological shifts. In contrast, native species may be at a competitive disadvantage when facing environmental stressors. Thus, intraspecific genetic diversity may have a key role in predicting the species potential dominance under unstable environmental regimes.

Metabarcoding using hypervariable markers constitutes an excellent tool not only for the study of marine communities at a given time point, but also to enable long-term monitoring studies of both biodiversity and genetic diversity. Implementing metabarcoding-based monitoring programs would allow the assessment of population status and responses to increasingly more frequent major climatic events (Berry et al., 2019). Such studies can provide an empirical framework for testing and refining ecological theories; traditionally developed in conceptual contexts or based on non-genetic information. The results obtained here allow the inference of ecological and evolutionary patterns that pave the way for future metagenomic and metatranscriptomic studies aimed at exploring the adaptive strategies observed in non-indigenous species.

## 5 Conclusions

Our results suggest that port communities are shaped by a complex interplay between abiotic factors, such as temperature and seasonality, and biotic factors, including competition and interspecific interactions, punctuated by alterations caused by extreme events. In our case, the exceptionally strong Storm Gloria acted upon an already pronounced seasonal dynamic, triggering a sudden reorganisation of biodiversity characterised by the arrival of new species and a temporary reduction in the relative abundance of non-indigenous species (NIS). The initial increase in species richness was followed by a progressive decline, although assessing community stability is challenging due to the absence of a robust pre-storm baseline and the influence of recent marine heatwaves recorded in the Mediterranean. Moreover, community structure showed increasing divergence over time, indicating structural shifts and a gradual evolution in species composition.

This study provides a detailed insight into the dynamics of NIS and their influence on port communities over a five-year period. Despite their lower species richness, NIS displayed high abundances and a notable ability to adapt to disturbed environments, reflecting their competitive advantage in these habitats. Our observations confirm that NIS exhibit lower compositional (match index) and genetic (D) differentiation over time than native species, revealing a higher structural and genetic homogenisation, likely facilitated by gene flow associated with maritime transport. At the same time, NIS showed higher haplotypic diversity than native species, indicating greater genetic variability and adaptive potential, traits that likely enhance their persistence and colonisation success in dynamic and environmentally stressed habitats.

Furthermore, species persisting across multiple years exhibited significantly higher genetic diversity, particularly among NIS, reinforcing the idea that genetic diversity acts as an insurance mechanism against environmental disturbances, consistent with the “insurance hypothesis”. These patterns highlight adaptive strategies that underpin the long-term success of NIS in port environments.

Overall, this long-term monitoring study improves our understanding of the role of NIS in shaping the structure and dynamics of marine communities, showing their capacity to maintain high connectivity and genetic diversity despite their low richness. It also demonstrates the value of metabarcoding using hypervariable markers as a tool to simultaneously assess biodiversity and genetic diversity in temporal studies, which are essential to detect relevant ecological changes and anticipate responses to climatic and biological disturbances. High-resolution temporal monitoring is crucial to guide management strategies, design effective conservation policies, and strengthen the resilience of marine ecosystems in the current context of accelerating biological invasions and climate change.

## Acknowledgments

The Department of Territory of the Catalan Government (“Ports de la Generalitat”) granted the permissions to work in the harbour surveyed. This research was funded by projects MARGECH (PID2020-118550RB) and BlueDNA (PID2023-146307OB) funded by the Spanish Ministry of Science, Innovation, and Universities (MICIU/AEI/10.13039/501100011033) and by ERDF/EU. J.Z. was funded by grant PRE2021-097703 (MICIU/AEI/10.13039/501100011033 and ESF+). Partial funding was also provided by the European Union (GA#101059915 - BIOcean5D, views and opinions expressed are those of the authors only and do not necessarily reflect those of the European Union. Neither the European Union nor the granting authority can be held responsible for them). AA was funded by the European Union’s Recovery and Resilience Facility-Next Generation within Investment 4 of Component 19 of the Recovery, Transformation and Resilience Plan. We also thank Mireia G. Mingote for helping on the first year of sampling.

## Disclosure

Benefit Sharing Statement: Benefits from this research accrue from the sharing of our data and results on public databases as described above.

## Author Contributions

The study was conceptualised by all authors. Project administration was managed by Owen Wangensteen, Marc Rius, and Xavier Turon. Funding acquisition was the responsibility of Owen Wangensteen, Marc Rius and Xavier Turon. Supervision and validation were carried out by Owen Wangensteen, Marc Rius, and Xavier Turon. Fieldwork was performed by Jesús Zarcero, Adrià Antich, Marc Rius, and Xavier Turon. Laboratory work was done by Jesús Zarcero and Adrià Antich. Data analyses were conducted by Jesús Zarcero and Xavier Turon. Software writing was undertaken by Jesús Zarcero and Xavier Turon. The original manuscript was drafted by Jesús Zarcero. Manuscript review and editing were undertaken by all authors.

## Declaration of Interests

The authors state that the present work was carried out in the absence of any commercial or financial relationships that might be interpreted as a potential conflict of interest.The authors state that the present work was carried out in the absence of any commercial or financial relationships that might be interpreted as a potential conflict of interest.

---

**Fig. S1.** Graph showing water temperature values at Blanes Port during the study period (February 2019 to February 2024).

**Fig. S2.** Rarefaction curves showing the number of MOTUs recovered at increasing read depths (up to 100,000 reads for clarity) in samples collected over the five-year study period.

**Fig. S3.** MOTU accumulation curves as a function of increasing sample number. Curves are shown by area and correspond to all detected metazoan MOTUs.

**Fig. S4.** (A) Bar charts showing the relative proportions of reads for all MOTUs from various phyla analysed at each station throughout the five-year study period. (B) Bar charts showing the relative proportions of reads for NIS MOTUs from various phyla analysed at each station throughout the five-year study period. (C) Bar charts showing the relative proportions of reads for NAT MOTUs from various phyla analysed at each station throughout the five-year study period.

**Fig. S5.** (A) Upset plots of shared MOTUs across years. (B) Upset plots of shared NIS MOTUs across years. (C) Upset plots of shared native MOTUs across years.

**Fig. S6.** Box plots showing Shannon diversity values for each year of the study: whole dataset (A), NIS dataset (B), and NAT dataset (C). Horizontal lines indicate medians, boxes show the first and third quartiles, whiskers indicate the 10th and 90th percentiles, and outliers are shown as dots.

**Fig. S7.** Box plots of MOTU richness values for each month of the study: whole dataset (A), NIS dataset (B), and NAT dataset (C).

**Fig. S8.** Box plots showing Shannon diversity values for each month of the study: whole dataset (A), NIS dataset (B), and NAT dataset (C).

**Table S1.** Table of the 2,225 metazoan MOTUs identified in the analyses. For each MOTU, the following information is provided: ID, PID, name, taxonomic rank assigned, taxonomic assignments at each level, total read abundance, abundance in each sample (see “Sample metadata” sheet), representative sequence, and category: NIS, NAT, or COMM (remaining community).

**Table S2.** Table of the 20,351 metazoan ESVs used in the analyses. For each ESV, the following information is provided: ID, total read abundance, MOTU assignment, abundance in each sample (see “Sample metadata” sheet), and sequence.

**Table S3.** (A) Results of the PERMANOVA tests and pairwise comparisons on the Bray-Curtis dissimilarity matrix for all MOTUs. (B) Results of the PERMANOVA tests and pairwise comparisons on the Bray-Curtis dissimilarity matrix for NIS MOTUs. (C) Results of the PERMANOVA tests and pairwise comparisons on the Bray-Curtis dissimilarity matrix for NAT MOTUs. (D) Table with Normality, Homogeneity of Variance and Pairwise tests for the Shannon diversity and MOTU richness variables for all MOTUs. (E) Table with Normality, Homogeneity of Variance and Pairwise tests for the Shannon diversity and MOTU richness variables for NIS MOTUs. (F) Table with Normality, Homogeneity of Variance and Pairwise tests for the Shannon diversity and MOTU richness variables for NAT MOTUs. (G) Forward stepwise regression analyses performed on NIS and NAT MOTUs. The cells indicate the decrease (in percentage) in AIC obtained adding each of the variables. The variable with higher decrease is selected (underlined) at each step. The procedure continues until there is no further decrease in AIC (positive values) adding more variables or all variables have been included. The adjusted r-squared obtained at each step is indicated *Supplementary material available at:* https://github.com/jesuszarcero/TEMP_supplementary_data/

## Notes

https://github.com/jesuszarcero/TEMP_supplementary_data

### Competing Interest Statement

The authors have declared no competing interest.

